# Temporal Changes in The Proanthocyanidins to Anthocyanins Ratio During Dormancy Associate with Bloom Time Variations in Peach

**DOI:** 10.1101/2023.06.13.544853

**Authors:** Protiva Rani Das, Md Tabibul Islam, Jianyang Liu, Zongrang Liu, Chris Dardick, Sherif M. Sherif

## Abstract

This study provides a thorough exploration of the mechanisms regulating the onset of flowering in peach trees, a process principally governed by bud-dormancy. We applied untargeted metabolomics combined with a comprehensive series of molecular and biochemical experiments to scrutinize the variations in bloom times among different peach cultivars. The impact of exogenous chemical stimuli, specifically ethephon (ET) and abscisic acid (ABA), on bloom times was also evaluated. Our study revealed that the ET-induced delay in bloom time was associated with higher levels of proanthocyanidin (PA) compared to anthocyanins (ACNs) during endodormancy. Furthermore, fluctuations in the PA/ACNs ratio during dormancy demonstrated a strong correlation with the chill requirements and bloom dates of 12 distinct peach genotypes. The research further uncovers the crucial role of ABA in regulating the biosynthesis of PAs and ACNs during peach tree dormancy. Intriguingly, the exogenous application of ABA during endodormancy resulted in a reduction of PA content, leading to an earlier bloom time. We also observed variations in DAM gene expression between early- and late-blooming cultivars. The late-blooming cultivars exhibited higher transcript levels of DAM genes, elevated PA levels, and lower ABA levels compared to their early-blooming counterparts. Importantly, our study proposes PAs and ACNs as quantitative marker metabolites for endo- and ecodormancy phases. This innovative finding paves the way for developing more accurate chill and heat requirement models, thereby enabling a more precise understanding and projection of the impacts of global climate change on the phenology of tree fruit species.

## Introduction

The onset of flowering in peach and other deciduous trees is largely regulated by bud dormancy. Dormancy can be defined as a period of growth inactivity during fall and winter that safeguards the vegetative and flower meristems from the detrimental effects of low temperatures and dehydration (Romeu et al. 2014). Three different forms of bud dormancy are observed in deciduous perennial trees, including paradormancy, endodormancy, and ecodormancy (Lang 1987). Paradormancy, exemplified by apical dominance, occurs early in the summer after terminal bud formation, in which the terminal bud curbs the growth of lateral buds. In contrast, endodormancy involves mechanisms intrinsic to the bud, and buds remain dormant despite the removal of terminal buds or defoliation. The resumption of active growth following endodormancy requires the fulfillment of specific chilling requirements (CRs) (Lang et al. 1987). CR is known to be genotype-dependent and highly diverse across species and varieties. For instance, some peach varieties require as high as 1000 chilling hours (CH), while others require only 50 CH (Layne and Bassi 2008). After the fulfillment of CR, floral buds undergo another phase of dormancy known as ecodormancy. Ecodormancy is governed by environmental factors such as cold and drought that activate signals and pathways, hindering bud growth. During this stage, buds necessitate a period of warm temperatures called heat requirements (HRs) before they can progress to bud break and blooming (Guo et al. 2014). Temperature-based models such as chilling hours (CH) and growing degree hours (GDH) (Weinberger 1950; Richardson et al. 1975) are the primary means of describing these intricate thermoregulatory dormancy cycles.

Bloom time in *Prunus* is a quantitative trait exhibiting high heritability, and determined, at least in part, by the genotype’s CR and HR (Layne and Bassi 2008). Multiple genetic studies have identified several quantitative trait loci (QTLs) that control CRs and flowering time in peach, indicating that dormancy-associated MADS-BOX (*DAM*) genes play a significant role in regulating dormancy onset and exit (Zhu et al. 2020). For instance, a peach mutant lacking *DAM* genes, also known as *EVERGROWING* (EVG), does not enter a dormant state (Zhu et al. 2020). The plant hormone abscisic acid (ABA), well-known for its growth suppression effects, has been a central focus in bud dormancy research. Studies on peach (*Prunus persica*), pear (*Pyrus pyrifolia*), sweet cherry (*Prunus avium*), and grapevine (*Vitis vinifera*) have demonstrated that ABA levels significantly increase during dormancy, followed by a decline concurrent with chilling accumulation (Tuan et al. 2017; Wang et al. 2015; Zheng et al. 2015). Moreover, the interaction between ABA and *DAM* genes has been established, providing a direct link between genetic and hormonal regulation of the endodormancy period (Tuan et al. 2017; Singh et al. 2018). Several studies have also reported notable associations between dormancy progression and release, and reactive oxygen species (ROS) production/scavenging (Islam et al. 2021; Perez et al. 2021), carbohydrate metabolism (Anderson et al. 2005), and flavonoid pathways (Conrad et al. 2019; Guillamon et al. 2020). However, the complex interplay among these biochemical pathways and the central dormancy circuit governed by *DAM* and ABA remains unclear. Deciphering the biochemical basis of flowering time regulation is expected to lead to the development of climate-resilient germplasm, frost-avoidance strategies, and ultimately, sustainable agriculture.

Spring frost damage is a significant economic concern in tree-fruit industries that occurs when ambient temperatures drop below the developmental stage-specific critical threshold, leading to the death of newly developed flowers and fruitlets and subsequently poor crop yields. However, despite the intuitive belief that climate warming would reduce frost damage, it is more likely to exacerbate the situation. Liu et al. (2018) showed that although climate warming reduces the total number of frost days worldwide, the number of frost days during the growing season increased in Europe and central North America over the last 30 years. This increase is attributed to a shift in the timing of the growing season, with earlier onset of spring leading to greater exposure to late-season frost events. Another negative effect of climate change is the alteration of tree phenology, which predicts that several phonologically responsive species in the US and Europe, including fruit trees, will experience more frost damage in the future (Ma et al. 2018). In Europe, for example, increasing spring temperatures over the last three to four decades have advanced apple blooming by an average of 2-3 days per decade (Vitasse et al. 2018a), resulting in an increased risk of frost, particularly for apples growing at higher elevations (Vitasse et al. 2018b). Traditional frost management techniques, such as microclimate-alteration, are costly, time-consuming, unpredictable, and environmentally unsustainable (Liu and Sherif 2019). However, the fall application of ethephon, an ethylene-based plant-growth regulator, shows promise in delaying bloom time in several stone fruits, including peach (Pahwa and Gahi 2015; Liu et al. 2021). Our previous studies have shown that ET delays bloom time in peaches by modulating CR, HR, cold hardiness, hormonal balance, reactive oxygen species (ROS) levels, antioxidant enzyme activity, carbohydrate metabolism, cell division, and intercellular transport (Liu et al. 2021; Islam et al. 2021; Liu et al. 2022).

In this study, we endeavored to elucidate the complex biochemical mechanisms that regulate bud dormancy and bloom time in peach. We employed an experimental model system utilizing ET-mediated bloom delay to decipher the underlying metabolic pathways that trigger dormancy release and bloom time. Floral buds from ET-treated and control peach trees underwent untargeted metabolomic profiling to identify critical marker metabolites and complex biochemical pathways underlying variations in dormancy duration and bloom time. Furthermore, we examined several peach cultivars exhibiting natural variations in CRs and bloom time to validate our ET system findings. Our results suggest that ET treatment upregulates PAs levels during endodormancy, suppressing ACNs biosynthesis during ecodormancy, and resulting in bloom delay. We also proposed a novel model where the PAs/ACNs ratio controls peach’s bloom time via a temperature, *DAM* genes, and ABA regulated pathway.

## Materials and methods

### StiPlant materials

In this study, we examined 12 peach cultivars with a wide range of chilling hours and diverse bloom dates. ‘Redhaven’ was planted in 2012, while ‘Sunhigh’, ‘Rich May’, and ‘Victoria’ were planted in 2017 at the Alson H. Smith Jr. Agricultural Research and Extension Center (AHS Jr. AREC) in Winchester, VA, USA (39.11, -78.28). These cultivars were grafted onto ‘Lovell’ rootstock. Eight additional cultivars and accessions, including ‘John Boy’, Y142-75, ‘Crimson Rocket’, ‘Contender’, A72, ‘Bailey’, ‘Gloria’, and ‘KV021779’, were planted at the USDA-ARS Appalachian Fruit Research Station in Kearneysville, WV (25430), USA (39.36, 77.86). ‘Contender’, Y142-75, and ‘Bailey’ were grafted onto ‘Lovell’ rootstock and planted between 2002 and 2006. ‘John Boy’, ‘A72’, and ‘Gloria’ were grafted onto ‘Bailey’ rootstock and planted between 2005 and 2016. ‘Crimson Rocket’ was planted in 2016 and grafted onto ‘Controller 5’ rootstock, while ‘KV021779’ was planted in 2003 and grafted onto its own roots. The agronomic characteristics of each peach genotype used in this study are presented in Table S1, and schematic diagrams depicting orchard layout, satellite views, and orchard locations are provided in Fig. S1 - S4.

It should be noted that ‘Redhaven’ peach trees at the AHS Jr. AREC served as the primary plant material, which underwent ethephon treatment and subsequent metabolomic analysis. A similar ‘Redhaven’ peach block at the same location was also utilized for ABA experiments. The other 11 cultivars were employed to evaluate the relationship between their dormancy and flowering characteristics and the proanthocyanidin (PAs) and anthocyanin (ACNs) ratio during dormancy. The following sections will provide detailed descriptions of the orchard layout and experimental design for each of these experiments.

### Treatments and experimental design

The foundational experiment for the present study, which compared ethephon (ET) treated and control (untreated) ‘Redhaven’ peach trees, has been previously described in the authors’ publications (Liu et al. 2021; Islam et al. 2022; Liu et al. 2022). This research utilized the same experiment and biological samples to investigate the effects of ethephon treatments on various factors such as chilling and heat requirements, cold hardiness, plant hormone levels, reactive oxygen species, antioxidation activities, carbohydrate levels, and transcriptomic changes. The current study examines alterations at the metabolomic level using a subset of the same biological samples collected at specific intervals of chilling hours (CH: 200, 400, 600, 800, 1000) and growing degree hours (GDH: 1000, 3000). It should be noted that only five of the seven time points (200 CH, 600 CH, 1000 CH, 1000 GDH, and 3000 GDH) were used for whole metabolome profiling, but all time points were utilized for the quantification of individual metabolites and related gene expression analyses (Fig. S1).

The ET experiment was conducted at the AHS Jr. AREC in Winchester, VA, United States, utilizing seven-year-old ‘Redhaven’ peach trees grafted to ‘Lovell’ rootstock and arranged in three adjacent rows (Fig. S2). Due to a slight elevation difference among rows, ET-treated and control trees were organized following a Randomized Complete Block Design (RCBD), with each row serving as a block (Fig. S2). In each block, two groups of two adjacent trees were randomly assigned to either receive ethephon treatment or remain as controls, with two buffer trees placed between the treated and control trees to prevent spray drift. Motivate® (Fine American Inc., Walnut Creek, CA, United States), containing 21% ethephon (2 chloroethylphosphonic acid), was used in this study. A mixture of ethephon (500 ppm) and nonionic surfactant Regulaid (250 ppm, Kalo Inc., Overland Park, KS, United States) was applied using an air-blast sprayer at 50% leaf fall (October 24, 2019). From each group, one tree was used for collecting floral bud samples, while the other tree was employed for branch collection, chilling and heat accumulation assessment, and monitoring flower progression and fruit set data. At each bud dormancy time point, a sufficient number of floral buds were randomly collected from one-year-old branches throughout the tree (biological replicate), pooled in a 15 mL tube, immediately frozen in liquid nitrogen, and stored at -80°C for subsequent biochemical and gene expression analyses (Fig. S1). Schematic illustrations of the ET experiment layout, floral bud sample collection, and treatment and replicate distribution in the peach orchard are provided in Fig. S1 and Fig. S2.

For the exogenous abscisic acid (ABA) treatments on peach trees, ABA (ProTone, Valent BioSciences, Inc., Libertyville, IL, USA) at concentrations of 100 and 500 mg/L (pH 5.0) was applied to seven-year-old ‘Redhaven’ peach trees located at the AHS Jr. AREC in a separate orchard consisting of seven adjacent rows with no significant elevation differences among rows. As a result, both untreated (control) and ABA-treated peach trees were arranged randomly in a single row following the Completely Randomized Design (CRD), with three biological replicates (one tree per replicate) per treatment (Fig. S3). ABA applications were conducted on December 13, 2021 (500 chilling hours) using a backpack sprayer after mixing with Regulaid (250 mg/L). Similar to the ethephon (ET) experiments, floral buds were randomly collected from one-year-old branches throughout the tree, combined in a 15 mL tube, immediately frozen in liquid nitrogen, and stored at -80°C for later biochemical and gene expression analyses.

To investigate the correlation between PAs and ACNs levels and the dormancy stage among peach genotypes, floral buds were collected from the 12 peach genotypes with varying chilling requirements and bloom times as described in the previous section. Peach cultivars including their geographical locations and the specific analyses performed are presented in Table S2. Floral bud samples were gathered from each peach genotype at 800 CH, 1000 CH, 1000 GDH, and 3000 GDH between January 14 and March 21, 2022, using ‘Redhaven’ as the standard cultivar for determining sample collection timing. For each peach cultivar/genotype accession, three peach trees were used as biological replicates, and bud samples were collected from each tree and stored following the methods previously mentioned.

### Estimation of chilling and heat accumulation and evaluation of bloom progression

Chilling and heat accumulation were calculated using meteorological data obtained from each orchard. Data loggers (EasyLog, Lascar, Erie, PA, United States) recorded temperatures at 10-minute intervals, placed in enclosed wooden shelters at 1.2 meters above the ground. Chilling accumulation was estimated as chilling hours (CH) based on the 0–7.2 °C model by Weinberger (1950). Heat accumulation was calculated as growing degree hours (GDH) based on the 4.5–25°C model by Richardson et al. (1975). Dormancy release’s chilling requirement (CR) was determined using the method described by Liu et al. (2021), where 50% bud break was observed under forcing conditions. Bloom progression was evaluated using the method described by Liu et al. (2021). The bloom rate was calculated as the ratio of open blossoms to the initial number of buds per tree, and the flowering date (F50) was determined for each treatment when the blooming rate reached 50%.

### Untargeted metabolomics

For untargeted metabolomics study, peach floral buds (around 50 mg) were extracted with 800 μL of 80% MeOH according to a previously described method (Das et al. 2021). Separation was performed using Ultimate 3000LC combined with Q Exactive MS (Thermo) and screened with ESI-MS. The mobile phase was composed of solvent A (0.05% formic acid-water) and solvent B (acetonitrile) with a gradient elution for ESI+ and ESI– mode. The most significant metabolite’s MS/MS spectra were acquired and searched using the Human Metabolome Database (www.hmdb.ca). Data processing and statistical analysis was performed according to our previous study (Das et al. 2021). Raw data were acquired and aligned using the Compound Discover (3.0, Thermo Fisher Scientific) based on the *m/z* value and the retention time of the ion signals. Ions from ESI+ or ESI were merged and imported into the SIMCA-P program (version 14.1). For multivariate statistical analysis, MetaboAnalyst 5.0 online software (https://www.metaboanalyst.ca/MetaboAnalyst/home.xhtml) was used.

### Gene expression analyses

Total RNA extraction was performed according to the CTAB method (Liu et al. 2022). Extracted RNA samples were purified using EZRNA Clean-Up Plus DNase Kit (EZ BioResearch, St Louis, Missouri, USA). The cDNA was synthesized from 2 μg of DNase-treated RNA using a cDNA synthesis kit (Applied Biosystems, Foster City, USA) according to the manufacturer’s instructions. Quantitative real-time-PCR (qRT-PCR) was performed using CFX connect real-time detection System (Bio-Rad, Mississauga, ON, Canada), SsoFast™ EvaGreenR Supermix (Bio-Rad), and gene-specific primers (Table S3).

### Total proanthocyanidin (PAs) and anthocyanin (ACNs) content

Peach floral buds (around 200 mg) were extracted with 1 mL of 80% MeOH and kept in a shaker incubator for 4 h at room temperature. Total ACNs and PAs content were determined using the Anthocyanins and Proanthocyanidins Assay Kit (Cosmo Bio, Carlsbad, CA, USA) according to the manufacturer’s instructions. Total ACNs and PAs content were expressed as micrograms per gram of fresh weight (µg/g FW).

### HPLC analysis

Quantification of epicatechin, procyanidin B1, cyanidin-3-glucoside, and ABA of was performed on the 80% methanol extract using an LC-40D (Shimadzu; Kyoto, Japan) system equipped with the SPD-M40 photo diode array detector (Shimadzu; Kyoto, Japan) and a 5 μm premier C18 column (250×4.60 mm) (Shimadzu; Kyoto, Japan). Separation was carried out following a previously described method (Das and Eun 2018) with 0.1% orthophosphoric acid in water (V/V) (eluent A) and 0.1% orthophosphoric acid in methanol (V/V) (eluent B). Detection of epicatechin and procyanidin B1 was carried out at 280 nm, cyanidin-3-glucoside at 518 nm, and ABA at 254 nm wavelength. Each metabolite (from Sigma-Aldrich Co., St Louis, MO, USA) was used as the internal standard for the calibration curve and their content were expressed as micrograms per gram of fresh weight (µg/g FW).

### 4-dimethylaminocinnamaldehyde (DMAC) staining of peach floral buds

Longitudinal sections were prepared from floral buds collected at 3000 GDH using a razor blade, and immediately soaked in water for 2 h at room temperature. Afterward, bud sections were soaked in 0.01% DMAC solution in absolute ethanol for 30 min at room temperature, then thoroughly washed with 70% ethanol. Images were taken immediately at this stage for clear visualization of stains.

### Statistical analysis

Data for metabolite quantification and gene expression analyses were presented as mean ± SE (standard error). To determine significant differences between experimental groups, analysis of variance (ANOVA) was performed using SAS 9.1.3 software (Cary, North Carolina, USA). In cases where significant differences were detected, a post-hoc analysis using Duncan’s Multiple Range test was conducted to compare means, and differences were considered statistically significant when p < 0.05. To assess significant differences between two specific groups, the Tukey test was employed, with significance levels denoted as follows: * at p ≤ 0.05, ** p ≤ 0.01, and *** p ≤ 0.001. It is important to note that the Tukey test is a more conservative post-hoc test, which helps control the family-wise error rate when multiple pairwise comparisons are made. All figures were constructed using Prism statistical software (GraphPad Prism 9.3.1, 2365 Northside Dr. Suite 560, San Diego, CA 92108, USA).

## Results

### Metabolite Profiling Revealed Differential Metabolic Regulation During Endo- and Ecodormancy Periods in Peach Floral Buds Treated with Ethephon

The fall application of ethephon (ET) significantly delayed the bloom time by 8 days, extended the duration of endo- and ecodormancy periods, and increased the CRs and heat requirements in the ‘Redhaven’ cultivar (Fig. 1, A and B). In a previous study, we reported on the ET-mediated bloom-delay effect (Liu et al. 2021; Islam et al. 2021). In the present study, we aimed to evaluate the comprehensive metabolite profile of peach floral buds at various endodormancy periods [200, 600, and 1000 chilling hours (CH)] and ecodormancy periods [1000 and 3000 growing degree hours (GDH)] by utilizing ultra-performance liquid chromatography time-of-flight mass spectrometry (UPLC-TOF/MS) to assess the untreated control and ET-treated group. We identified a total of 419 metabolites, including 136 in electrospray ionization (ESI)+ mode, 31 in ESI mode, and 252 in both ESI+ and ESI− modes (Table S4). Interestingly, we observed that all metabolites were present in the untreated control and ET-treated buds throughout the dormancy periods, but their relative abundances differed between the treatment and dormancy stages. Orthogonal partial least-squares discriminant analysis (OPLS-DA, the overview is presented in Fig. S5) identified 69 and 50 candidate metabolites [variable important parameter (VIP) > 1.5], primarily attributed to the significant fluctuation between the untreated control and ET-treated buds during endodormancy (Fig. 1, C and E; Table S5) and ecodormancy periods (Fig. 1, D and F; Table S6), respectively. Epicatechin emerged as the top-positioned metabolite during the endodormancy period, while asparagine was predominant during the ecodormancy period. Furthermore, metabolite set-enrichment analysis (MSEA) showed a marked variation in chemical groups between the endo- and ecodormancy periods (Fig. 1G). Notably, we observed a higher percentage of flavonoids (3-fold higher) in the endodormancy period relative to the ecodormancy period, while organic and amino acids were 2-fold higher in the ecodormancy period relative to the endodormancy period. These findings highlight the differential metabolic regulation during endo- and ecodormancy periods.

**Figure 1.**
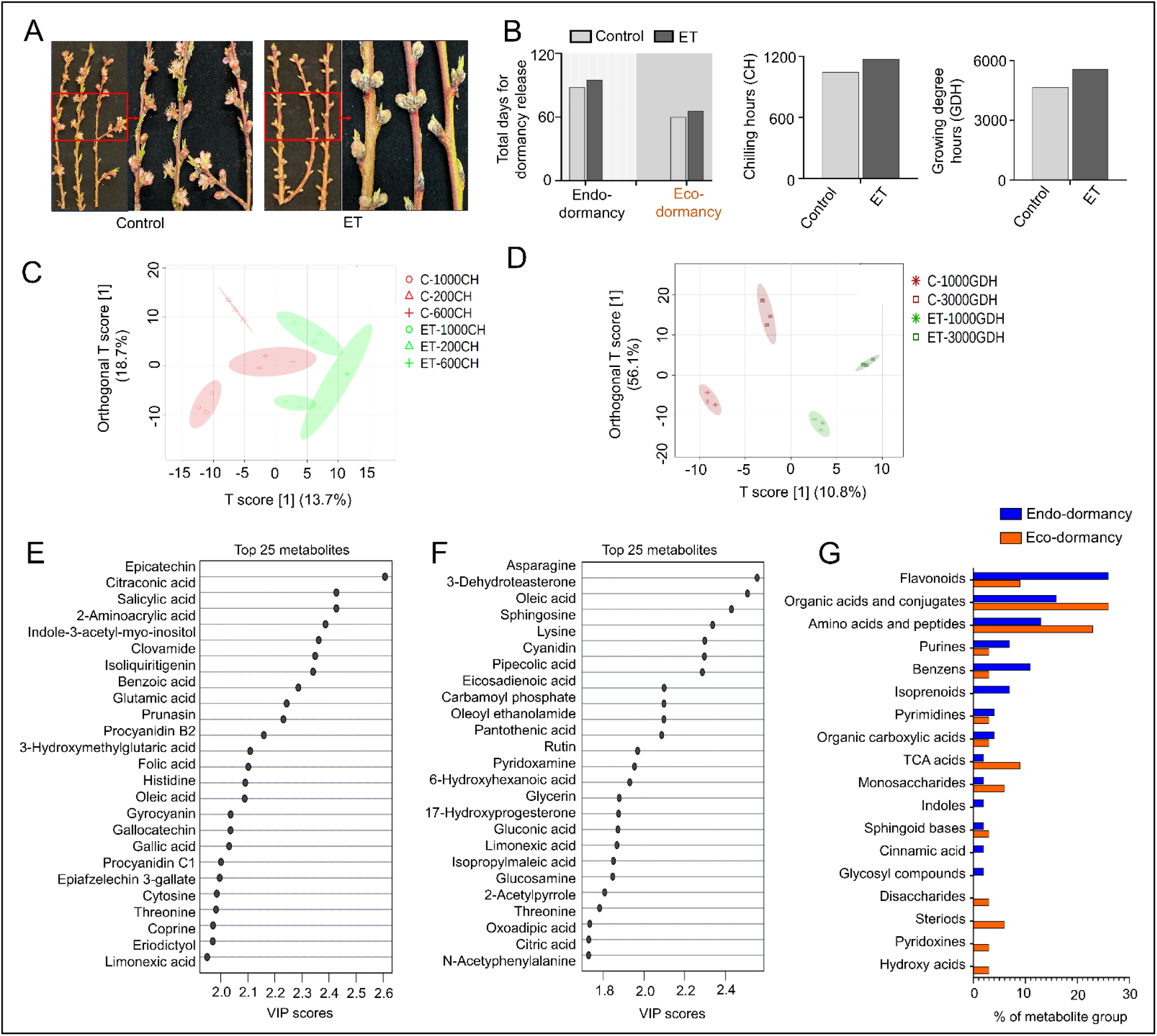
Multivariate statistical analysis determines candidate discriminant metabolites between untreated control and ethephon (ET)-treatment throughout the endo- and ecodormancy periods. **A**) ET treatment shows the bloom delay effect in ‘Redhaven’ cultivar. Branches are collected from the field to capture images, and an arrow points to zoom the image to make it more visible and apparent. Photos taken at full bloom time of untreated control (25^th^ March 2022). **B**) ET increased the total number of days for dormancy release and prolonged chilling and heat periods. **C, D**) Orthogonal partial least–squares–discriminant analysis (OPLS-DA) of 419 identified metabolites between untreated control and ET treatment during the endodormancy (200 CH, 600 CH, and 1000 CH) and ecodormancy (1000 GDH and 3000 GDH) periods, respectively. **E, F**) Top 25 variable importance in projection (VIP) scores metabolites that act as the significant discriminant between untreated control and ET treatment during endo- and ecodormancy periods, respectively. **G**) Over representative enrichment ratio analysis of candidate metabolites (VIP > 1.5) using metabolite set enrichment analysis (MSEA) during the endo- and ecodormancy periods. As all 419 metabolites were identified in both untreated control and ET treatment, MSEA represents the primary chemical classes of metabolites set with percentage (%). The MSEA analysis was performed using only the HMDB ID of identified compounds without their relative contents. In the 2D OPLS-DA score plot, the red color of the triangle, cross-shape, and oval shape represent untreated control at 200 CH, 600 CH, and 1000 CH; the green color of the triangle, cross-shape, and oval shape represents ET treatment at 200 CH, 600 CH, and 1000 CH of endodormancy periods. The red color of the star-shape and rectangle represents untreated control at 1000 GDH and 3000 GDH; the green color of the star-shape and rectangle represents ET treatment at 200 CH, 600 CH, and 1000 CH of ecodormancy periods. The blue and orange colors in the MSEA bar plot designate endodormancy and ecodormancy periods, respectively.

### Epicatechin as a Discriminatory Marker for Effects of ET Treatment at Each Endodormancy Period

To identify the marker metabolite associated with ET-mediated bloom delay, we performed multivariate statistical analyses of candidate metabolites (VIP > 1.5) during the endo- and ecodormancy periods (Fig. 2 and 3). Hierarchical clustering analysis (HCA) evaluated metabolite discrimination at each endodormancy period (Fig. 2A), showing contrasting patterns between the untreated control and ET-treated buds. A visual difference was observed between the untreated control and ET-treated buds at 1000 CH, the crucial endodormancy-release period. Pattern search plot analysis identified the top 20 marker metabolites at each endodormancy period, revealing significant upregulation of 10 positively correlated and downregulation of 10 negatively correlated metabolites in the ET-treated and untreated control buds, respectively (Fig. 2 B–D). Epicatechin emerged as the top-positioned, positively correlated marker metabolite in each endodormancy period, showing 1.2-, 1.2-, and 1.4-fold higher intensity levels relative to the untreated control at 200 CH, 600 CH, and 1000 CH, respectively (Table S7). Additionally, epicatechin showed a higher positive correlation with metabolites in a common metabolic pathway or with similar trends of variation (Fig. S6). The metabolic pathway activated during endodormancy periods, flavonoid biosynthesis, appeared to be the most significant (Fig. 2E; Table S8), with six marker metabolites involved in the pathway, including isoliquiritigenin, phloretin, naringenin, eriodyctiol, luteolin, and epicatechin (Fig. 2F, Fig. S7). The accumulation pattern of these marker flavonoids was higher during endodormancy periods and decreased dramatically after the endodormancy-release periods and ultimately bottomed during the later stage of ecodormancy (3000 GDH). These findings reveal flavonoid biosynthesis as a significant regulatory pathway for endodormancy maintenance and release and highlight epicatechin as a positive regulator in ET treatment during the endodormancy-release period.

**Figure 2.**
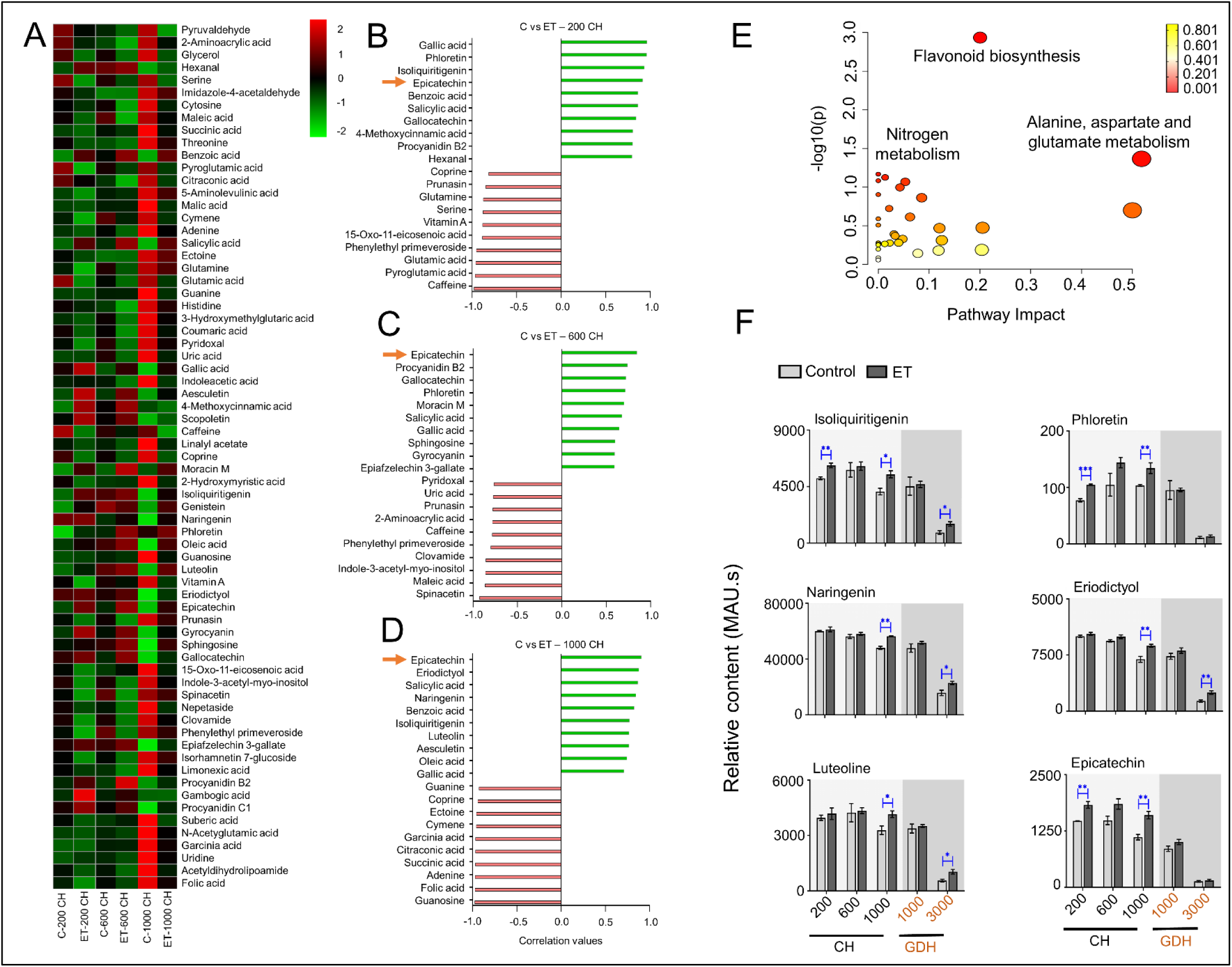
Epicatechin appeared as the top-position marker metabolite at each endodormancy period between untreated control and ET treatment. **A)** Hierarchical Clustering Heatmap (HCA) analysis of candidate metabolites (VIP > 1.5, a total of 69 metabolites) shows significant variations between untreated control and ET treatment during endodormancy. **B, C, D)** Pattern searching plot of top 20 metabolites with positive (green) and negative (red) correlations at 200 CH, 600 CH, and 1000 CH between untreated control and ET treatment, respectively. **E)** Pathway analysis shows flavonoid biosynthesis occurred as a dominant metabolic pathway at endodormancy stage. **F)** Relative abundances of six marker metabolites in the flavonoid pathway are significantly increased by ET treatment during endodormancy release periods. The metabolic pathways are presented according to the *p* values from the pathway enrichment analysis and pathway impact values from the pathway topology analysis. The influence of primary metabolic pathways is determined based on the *p-value* (< *0.05*) and FDR (< 1.0). Data represent the mean values ± SE (n = 3). * Significant at *p ≤ 0.05*, ** at *p ≤ 0.01*, and *** at *p ≤ 0.001* based on T-Test analysis between untreated control and ET treatment. CH: chilling hours during endodormancy periods and GDH: growing degree hours during ecodormancy periods.

**Figure 3.**
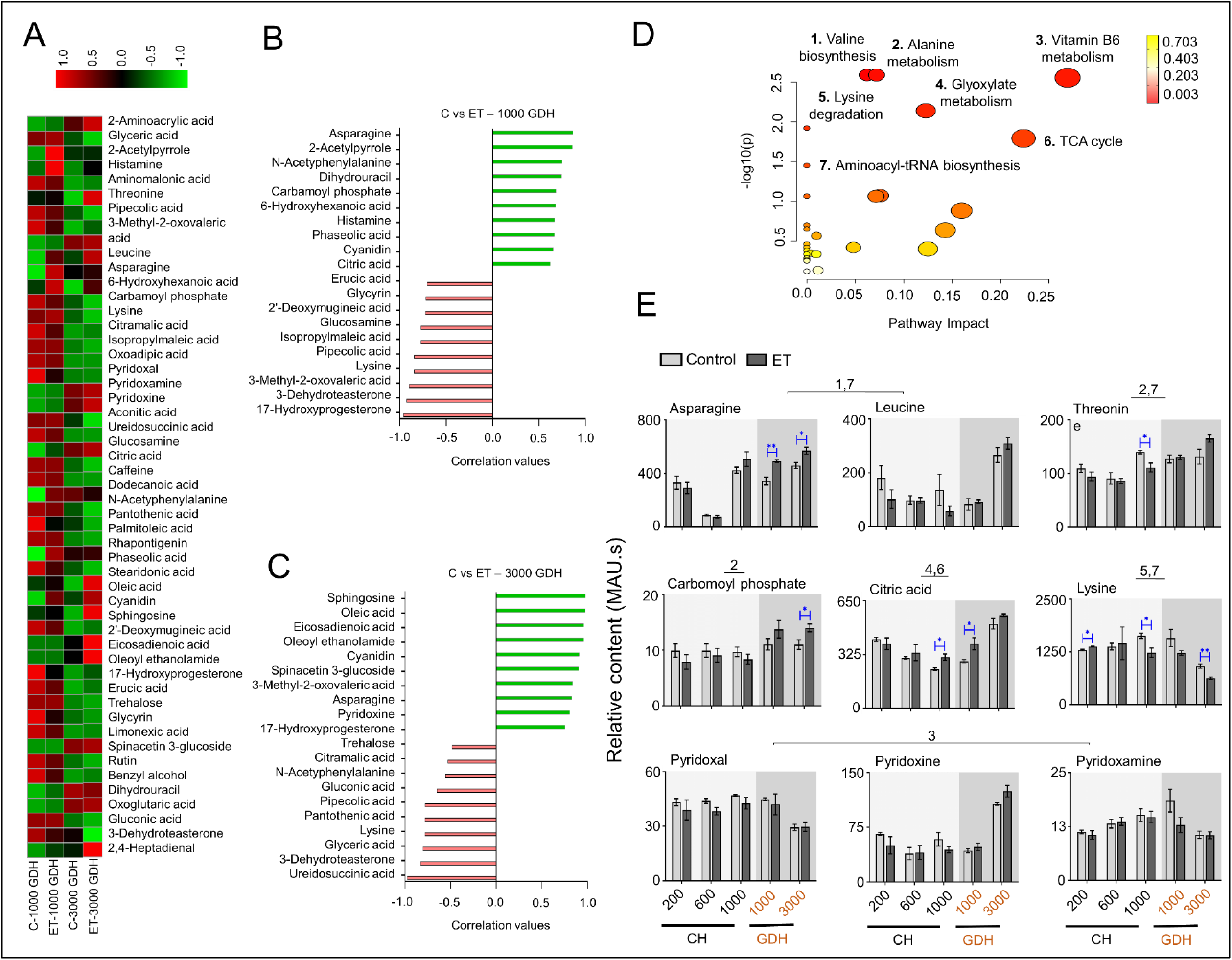
Amino acids are predominant during ecodormancy periods in untreated control and ET treatment. **A)** HCA analysis of candidate metabolites (VIP > 1.5, a total of 50 metabolites) shows significant variation between untreated control and ET treatment during ecodormancy. **B)** Pattern-searching plot of top 20 metabolites with positive (green) and negative (red) correlations at 1000 GDH and 3000 GDH between untreated control and ET treatment, respectively. **c** Pathway analysis shows seven amino acid-based biosynthesis is the significant metabolic pathway during ecodormancy periods. **D)** Relative abundances of nine marker metabolites in this pathway show different significance levels between untreated control and ET treatment throughout the dormancy periods. **1.** Valine, leucine, and isoleucine biosynthesis pathway; **2.** Alanine, aspartate and glutamate metabolism; **3.** Vitamin B6 metabolism; **4.** Glyoxylate and dicarboxylate metabolism; **5.** Lysine degradation; **6.** Citrate cycle; **7.** Aminoacyl-tRNA biosynthesis was found as the major metabolic pathway (*p > 0.05*) during ecodormancy periods. The metabolic pathways are presented according to the *p* values from the pathway enrichment analysis and pathway impact values from the pathway topology analysis. The influence of primary metabolic pathways is determined based on the *p-value* (< *0.05*) and FDR (< 1.0). Data represent the mean values ± SE (n = 3). * Significant at *p ≤ 0.05*, ** at *p ≤ 0.01*, and *** at *p ≤ 0.001* based on T-Test analysis between untreated control and ET treatment. CH: chilling hours during endodormancy periods and GDH: growing degree hours during ecodormancy periods.

### Amino Acid-Based Metabolites Showed Differential Variation During Ecodormancy Period in Buds Treated with Ethephon

The levels of amino acid-based metabolites varied significantly during the ecodormancy period, and unlike the endodormancy periods, the types of amino- and organic acid-based metabolites showed variation between the untreated control and ET-treated groups at each ecodormancy period (Fig. 3A). Asparagine and cyanidin were the common positively correlated metabolites in the ET-treated group, exhibiting 1.4-fold higher levels than those in the untreated control at 1000 GDH and 3000 GDH (Fig. 3, B and C; Table S9). Asparagine was also positively correlated with cyanidin, phaseolic acid, oleic acid, and histamine levels (Fig. S8). Seven amino acid biosynthesis pathways were found to be activated during the ecodormancy period (Fig. 3D; Table S10), and among the nine marker metabolites involved in these pathways, asparagine, citric acid, and carbamoyl phosphate showed significantly higher levels in the ET-treated group relative to the control (Fig. S9). In contrast, lysine was substantially higher in the untreated control during the ecodormancy period (Fig. 3E) These marker metabolites exhibited a decreasing trend as dormancy progressed, which was the opposite of the pattern observed during the endodormancy periods. Most of the metabolites, except lysine, pyridoxamine, and pyridoxal, peaked during the ecodormancy period. These findings suggest that metabolic activity of a particular amino acid corresponds to the progression of ecodormancy for growth and developmental signals.

### Ethephon Treatment Increased Flavonoid Biosynthesis and Proanthocyanidin (PAs) Content during Endodormancy Period

Because the length of the endodormancy period determines bloom time in peach (Lang et al. 1987), we analyzed the KEGG flavonoid pathway for the entire metabolome profile (419 metabolites) throughout the dormancy period to determine the significance of this pathway. Flavonoid biosynthesis showed significant activity throughout the dormancy period, generating a total of 15 flavonoids (Fig. 4A). Analysis of the relative normalized expression of genes related to this pathway showed significant variations between the untreated control and ET-treated groups throughout the dormancy period (Fig. S10). ET treatment significantly upregulated anthocyanidin synthase (*ANS*) and anthocyanidin reductase (*ANR*) expression during the endodormancy period, whereas it decreased dramatically with the progression of ecodormancy (Fig. 4, B and C). ET treatment significantly downregulated anthocyanidin 3-O-glucosyltransferase (*UFGT*) expression at the later stage of ecodormancy (3000 GDH), whereas it was significantly upregulated (by 12-fold) in the untreated control (Fig. 4D). Similarly, ET treatment significantly downregulated *3-GT* and *UGT78* expression at the later stage of ecodormancy (3000 GDH) (Fig. S10, J and K). ET treatment increased *MYBPA1* expression and PAs content during the endodormancy-release periods, whereas it decreased ACNs content along with downregulated MYB10 expression at the later stage of ecodormancy (3000 GDH) (Fig. 4, E–H). ET treatment also increased epicatechin and procyanidin B1 contents during the endodormancy period and decreased cyanidin-3-glucoside content during the ecodormancy period (Fig. 4, I–K). These findings demonstrate a clear transitional pattern of PAs to ACNs generation from the endo to the ecodormancy periods in both the untreated control and ET treated groups.

**Figure 4.**
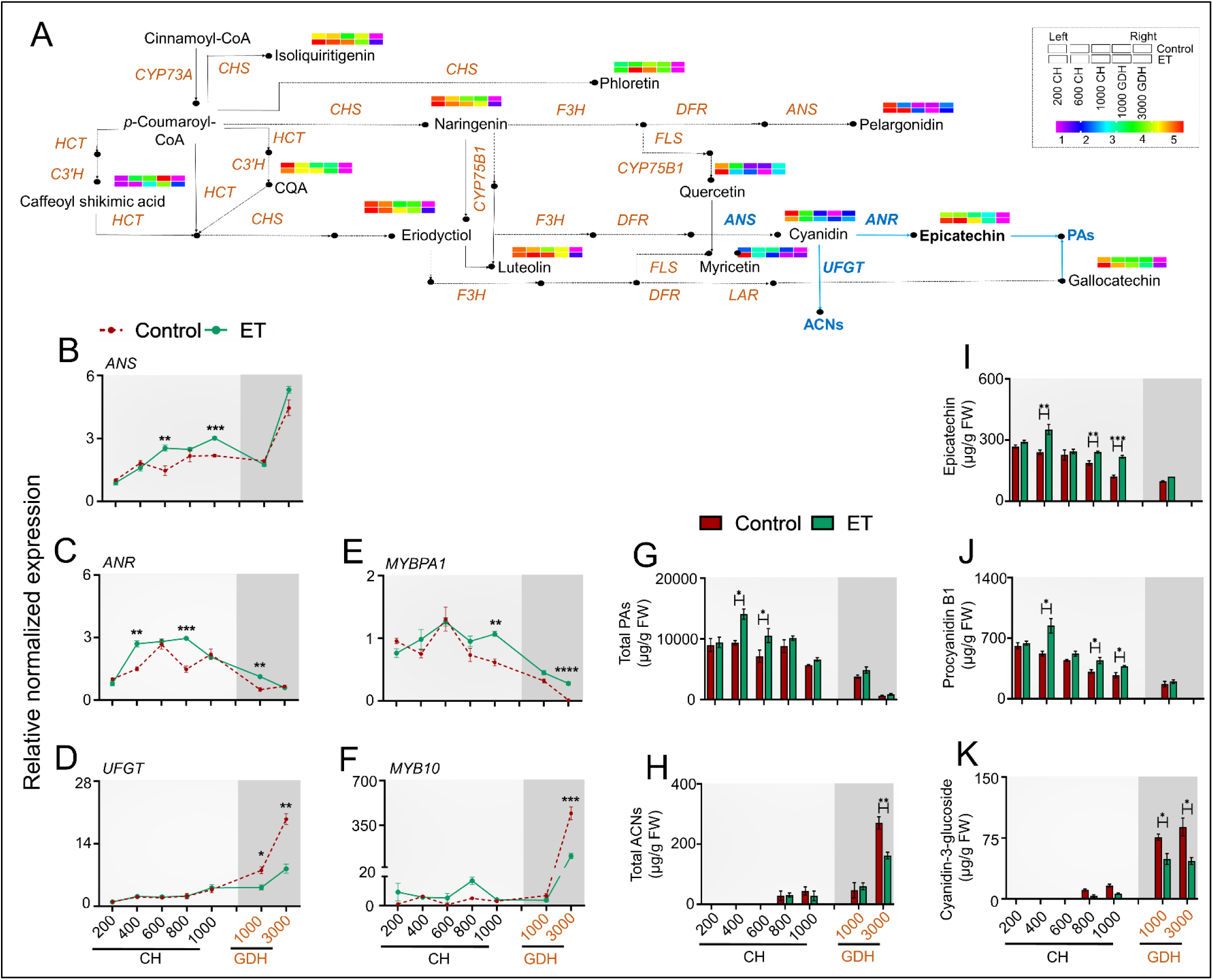
ET treatment increases proanthocyanidins (PAs) biosynthesis during endodormancy release but slow down anthocyanins (ACNs) biosynthesis during ecodormancy. **A)** Intensity levels of all metabolites (among whole 419 metabolites) involved in the flavonoid biosynthesis pathway along with highlighting corresponding genes throughout the dormancy periods. **B–D)** Relative expression of *ANS*, *ANR*, and *UFGT* in untreated control and ET treatment throughout the endodormancy (200 CH, 400 CH, 600 CH, 800 CH, and 1000 CH) and ecodormancy (1000 GDH and 3000 GDH) periods. **E**–**F)** Relative expression of *MYBPA1* and *MYB10* transcriptional factors in untreated control and ET treatment throughout the dormancy periods. **G–H**), Total PAs and ACNs content in untreated control and ET treatment throughout the dormancy periods. **I–K)** Quantification of epicatechin, procyanidin B1, and cyanidin-3-glucoside in untreated control and ET treatment throughout the dormancy periods. The expression of each gene was normalized to that of *β-Actin* and *Ubiquitin* and expressed relative to the untreated control (200 CH). Line graphs represent the relative gene expression levels, and the bar graph represents metabolite contents (Dark red and green color represents untreated control and ET treatment). Data represent the mean values ± SE (n = 3). * Significant at *p ≤ 0.05*, ** at *p ≤ 0.01*, and *** at *p ≤ 0.001* based on T-Test analysis between untreated control and ET treatment. *ANS*: anthocyanidin synthase, *ANR*: anthocyanidin reductase, *UFGT*: anthocyanidin 3-O-glucosyltransferase, *MYBPA1*: MYBPA1-like transcription factor, and *MYB10*: R2R3 MYB transcription factor. CH: chilling hours during endodormancy periods and GDH: growing degree hours during ecodormancy periods.

### PA/ACN Biosynthesis Pattern Correlated with Bloom Time in Peach Cultivars

To determine whether the biosynthetic pathway regulating PA/ACN levels acts as a marker metabolic pathway for endodormancy release and bloom time in peach, we quantified the PA and ACN contents in 12 peach cultivars with contrasting bloom times and CRs during two endodormancy (800 and 1000 CH) periods and two ecodormancy (1000 and 3000 GDH) periods (Table S1 and Table S2). Total PA content was significantly higher in the late-bloom cultivars during endodormancy, whereas total ACN content was substantially higher in the early-bloom cultivars during the ecodormancy period (Fig. 5, A and B). Additionally, staining the buds with DMAC dye showed a decline in the PA/ACN ratio at the later stage of ecodormancy (3000 GDH) that correlated with bloom time (Fig. 5, C and D). We investigated the integrated metabolite content and target gene expression in a late-bloom, ‘KV021779’ (CR 1308), and an early-bloom, ‘John Boy’ (CR 800), cultivars. The ‘KV021779’ cultivar contained higher contents of epicatechin and procyanidin B1 accompanied by upregulated *MYBPA1* and *ANR* expression during endodormancy periods, whereas the ‘John Boy’ cultivar showed higher cyanidin-3-glucoside content along with upregulated *MYB10*, *UFGT*, and *UGT78* expression during ecodormancy (Fig. 5, E–L; Fig. S11). These results provide further validation that genetically late-blooming cultivars contain higher levels of PA during endodormancy, which extends the dormancy-release period and results in later blooms.

**Figure 5.**
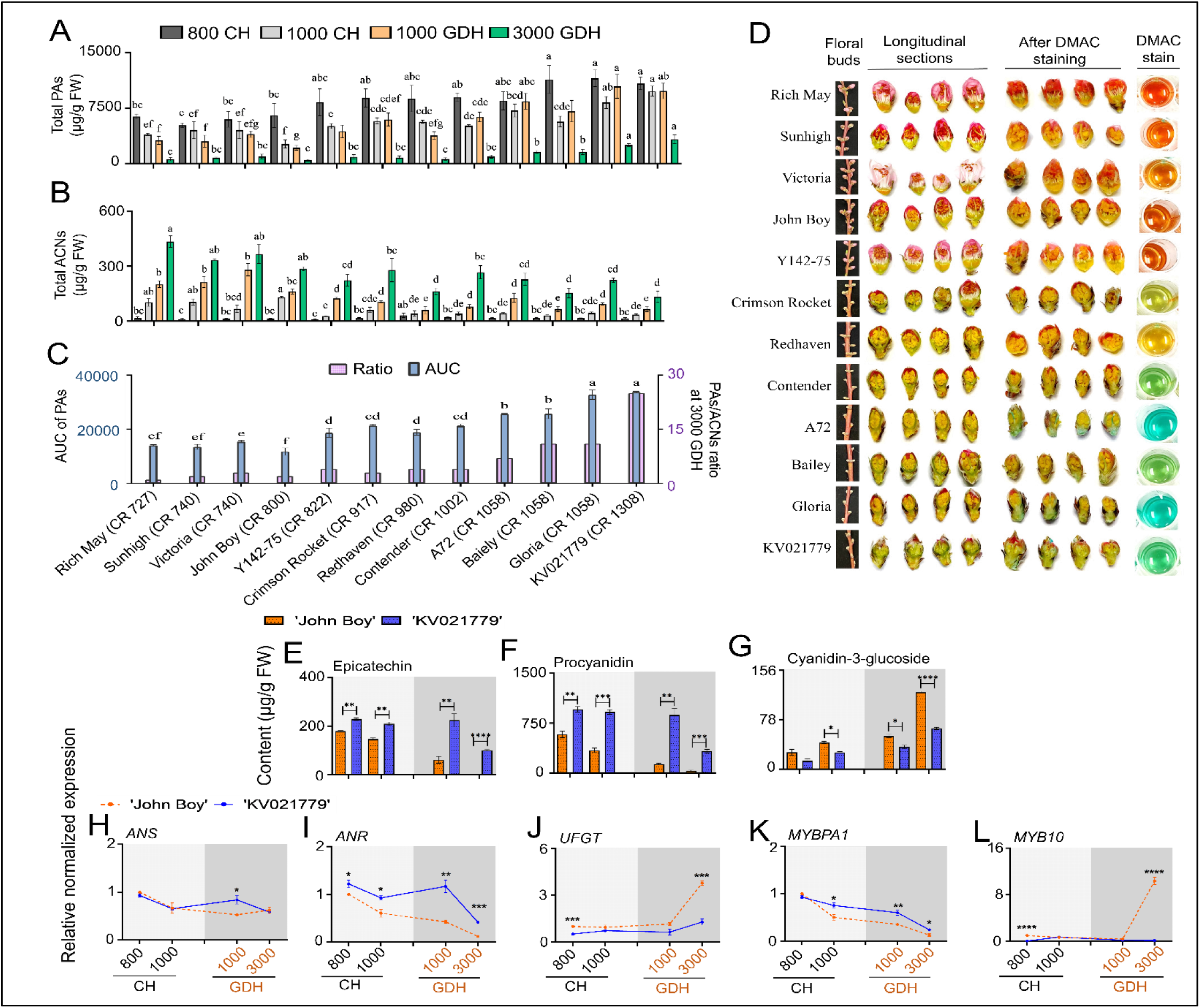
PAs/ACNs are the marker metabolites to determine early-bloom and late-bloom cultivars genetically. **A–B)** Total PAs and ACNs content in peach genotypes throughout the endodormancy (800 CH and 1000 CH) and ecodormancy (1000 GDH and 3000 GDH) periods. **C)** Area under the curve (AUC) of PAs throughout the endo- and ecodormancy periods (left Y-axis of the graph) and the ratio of PAs/ACNs at 3000 GDH of ecodormancy periods (right Y-axis of the graph). **D)** Morphological images and DMAC staining of peach floral buds at 3000 GDH of the ecodormancy period. **E–G)** Quantification of epicatechin, procyanidin B1, and cyanidin-3-glucoside in early-bloom ‘John Boy’ and late-bloom ‘KV021779’ cultivar throughout the endo- and ecodormancy periods. **H–L)** Relative expression of *ANS*, *ANR*, *UFGT, MYBPA1*, and *MYB10* in ‘John Boy’ and ‘KV021779’ throughout the endo- and ecodormancy periods. Colors representing CH and GDH of each genotype are indicated. The expression of each gene was normalized to that of *β-Actin* and *Ubiquitin* and expressed relative to the ‘John Boy’ (800 CH). a-d the different lowercase letters represent the significant differences among peach cultivars, according to Duncan’s multiple range test (*p > 0.05*). The bar graph represents metabolite contents, and line graphs represent the relative gene expression levels (Orange and blue color represent ‘John Boy’ and ‘KV021779’). Data represent the mean values ± SD (n = 3). * Significant at *p ≤ 0.05*, ** at *p ≤ 0.01*, and *** at *p ≤ 0.001* based on T-Test analysis between ‘John Boy’ and ‘KV021779’. *ANS*: anthocyanidin synthase, *ANR*: anthocyanidin reductase, *UFGT*: anthocyanidin 3-O-glucosyltransferase, *MYBPA1*: MYBPA1-like transcription factor, and *MYB10*: R2R3 MYB transcription factor. CH: chilling hours during endodormancy periods and GDH: growing degree hours during ecodormancy periods. Note that, based on the ‘Redhaven’ cultivar, 800 CH, 1000 CH, 1000 GDH, and 3000 GDH were determined for floral bud collection of all genotypes used in this study, where each genotype was at different developmental stages.

### ET Treatment Reduced ABA Levels and Affected Hormonal Regulation During Endodormancy Release

In order to investigate the hormonal regulatory role of ET treatment in the biosynthesis of PAs/ACNs and delay of bloom, we analyzed the untargeted metabolome profile (419 metabolites) for the presence of plant hormones. Our search identified three hormones, including salicylic acid and jasmonic acid, which were significantly more abundant in the ET-treated group compared to the untreated control. In contrast, ABA levels decreased significantly during the endodormancy-release time (1000 CH) (Fig. 6, A–C). Since the relative ABA content in the ET treated group showed an opposite regulatory pattern to PA levels during endodormancy release (Fig. 4), we further investigated the expression of three ABA-biosynthesis-related genes, *NCED1*, *NCED3*, and *NCED6*, throughout the dormancy period. Our results showed that ET treatment downregulated the expression of these genes during endodormancy (Fig. 6, D–F). Additionally, we evaluated ABA content and gene-expression levels in early-bloom ‘John Boy’ and late-bloom ‘KV021779’ peach cultivars throughout the dormancy periods. Interestingly, we found significantly higher ABA content in ‘John Boy’ relative to that in ‘KV021779’ throughout the dormancy period, with the expression of ABA-biosynthesis genes showing similar trends (Fig. 6, G–J). The negative correlation between decreasing ABA levels following ET treatment and late-bloom cultivars with higher PA levels (Fig. 4 and 5) strongly suggested that ABA plays a crucial role in regulating PA/ACN biosynthesis during endodormancy.

**Figure 6.**
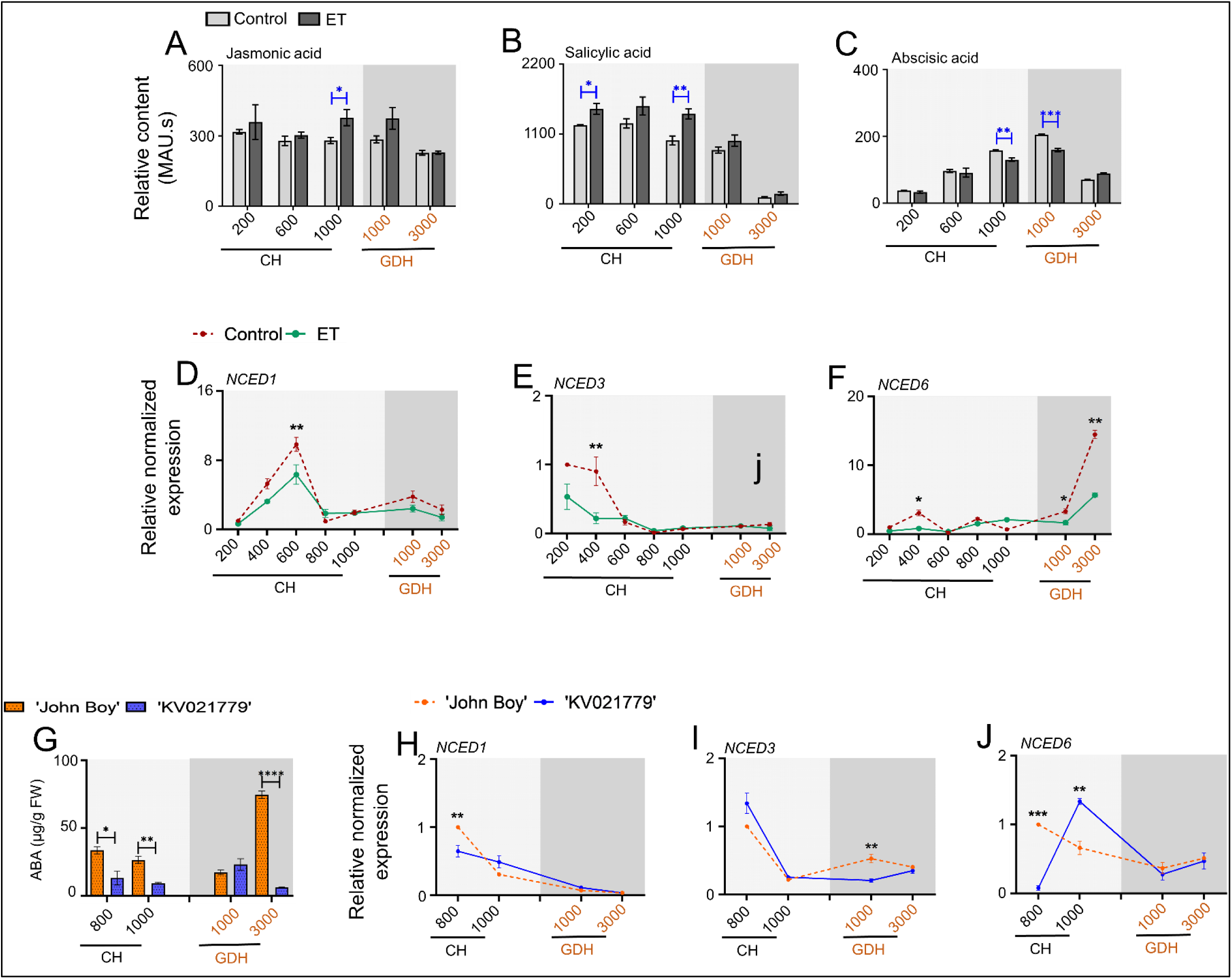
ABA levels significantly varied between untreated control vs. ET treatment and early-bloom and late-bloom cultivars during endodormancy release periods. **A–C)** Relative abundances (MAU.s) of JA, SA, and ABA in untreated control and ET treatment throughout the endo- and ecodormancy periods. **D–F)** Relative expression of ABA biosynthesis-related (*NCED1*, *NCED3*, and *NCED6*) genes in untreated control and ET treatment throughout the dormancy periods. **G)** ABA content (µg/g FW) in ‘John Boy’ and ‘KV021779’ throughout the endo- and ecodormancy periods. **H–J)** Relative expression of ABA biosynthesis-related (*NCED1*, *NCED3*, and *NCED6*) genes in ‘John Boy’ and ‘KV021779’ throughout the dormancy periods. The dark red and green color in the bar and line graphs represents untreated control and ET treatment. The expression of each gene was normalized to that of *β-Actin* and *Ubiquitin* and expressed relative to the untreated control (200 CH). The orange and blue in the bar and line graphs represent ‘John Boy’ and ‘KV021779’, respectively. The expression of each gene was normalized to that of *β-Actin* and *Ubiquitin* and expressed relative to the ‘John Boy’ (800 CH). Data represent the mean values ± SE (n = 3). * Significant at *p ≤ 0.05*, ** at *p ≤ 0.01*, and *** at *p ≤* 001 based on T-Test analysis. *NCED1*: 9-cis-epoxycarotenoid dioxygenase 1, *NCED3*: 9 cis-epoxycarotenoid dioxygenase 3, and *NCED6*: 9-cis-epoxycarotenoid dioxygenase 6. CH: chilling hours during endodormancy periods and GDH: growing degree hours during ecodormancy periods.

### Suppression of PAs Biosynthesis and Advancement of Bloom Time through Exogenous ABA Treatment during Endodormancy Period

As ABA is known to affect flavonoid biosynthesis (Guillamon et al. 2020; Zhao et al. 2019) and the current study showed a negative correlation with PA biosynthesis during endodormancy (Fig. 4–6), we investigated the potential link between ABA levels and PA biosynthesis. Applying exogenous ABA (100 and 500 mg/L) during endodormancy (500 CH) in a field trial resulted in an advanced bloom date by 2 to 3 days in the ‘Redhaven’ cultivar (Fig. 7, A–C). Notably, the application of 100 and 500 mg/L respectively significantly decreased PAs content and downregulated MYBPA1 and LAR expression in ABA-treated floral buds during endodormancy (Fig. 7, D–G). These findings suggested that ABA inhibited PA biosynthesis by downregulating ANR/LAR during endodormancy periods, leading to advanced bloom dates.

**Figure 7.**
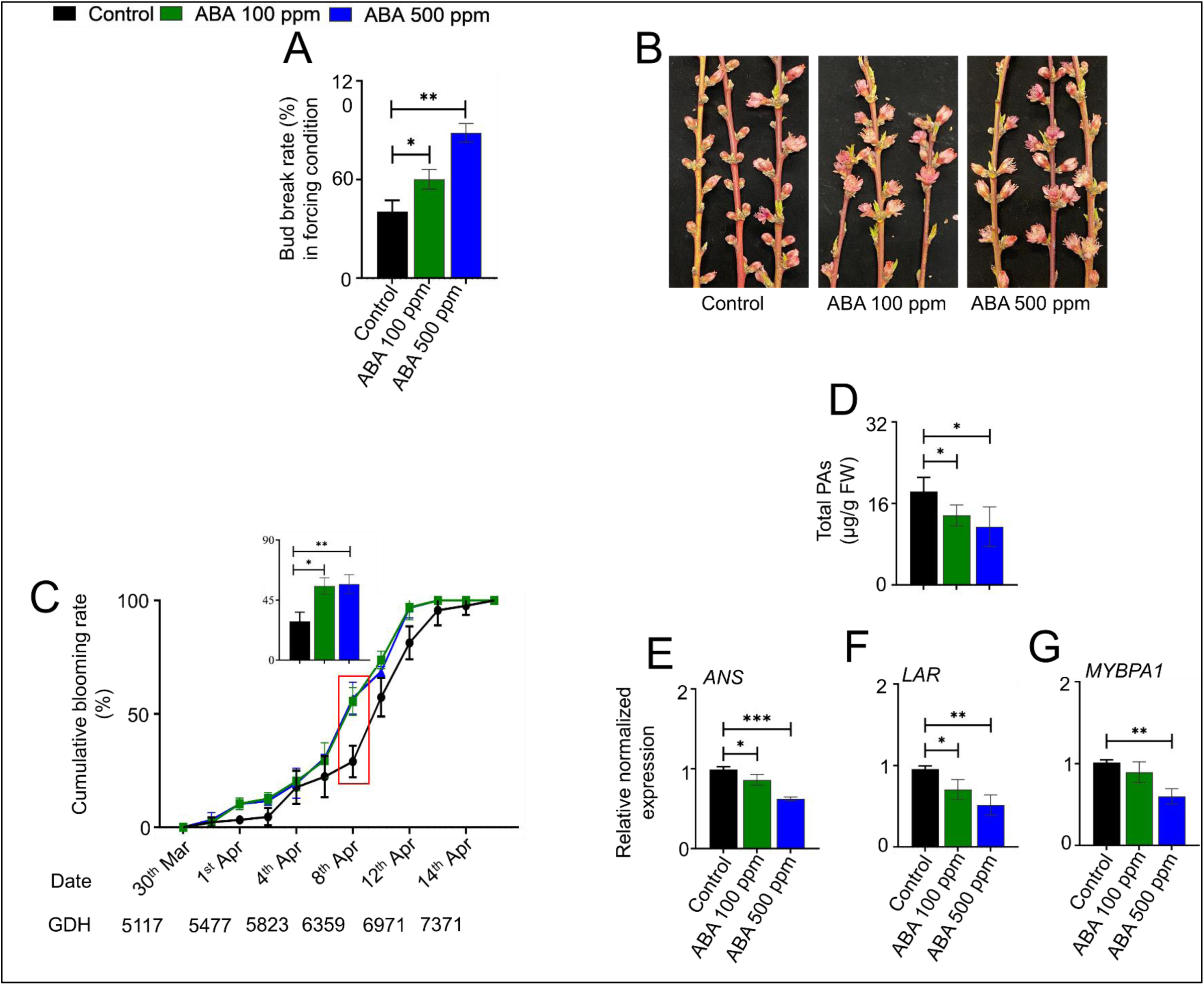
Exogenous ABA application during endodormancy periods decreases PAs biosynthesis in ‘Redhaven’. **A–C)** Exogenous ABA treatment at 500 CH shows the advanced blooming effect in ‘Redhaven’, photos taken at full bloom time of the ABA treatment. **B)** Blooming progression of ‘Redhaven’ buds as affected by ABA application and blooming rate calculated as the ratio of a number of flowers to the number of buds (around 100 floral buds) on tagged branches. **C)** Blooming progression in the growth chamber condition and blooming rate calculated as the ratio of the number of flowers to the number of buds on collected branches. **D)** Total PAs content (µg/g FW) in untreated control and ABA treatments after 48 hrs of application. **E)** Relative expression of *ANS*, *LAR*, and *MYBPA1* genes in untreated control and ABA treatments after 48 hrs of application. Black, green, and blue color represent untreated control, ABA 100 mg/L, and ABA 500 mg/L. The expression of each gene was normalized to that of *β-Actin* and *Ubiquitin* and expressed relative to the untreated control (0 hr). Data represent the mean values ± SE (n = 3). * Significant at *p ≤ 0.05*, ** at *p ≤ 0.01*, and *** at *p ≤ 0.001* based on T-Test analysis. *ANS*: anthocyanidin synthase, *LAR*: leucoanthocyanidin reductase/leucocyanidin reductase, and *MYBPA1*: MYBPA1-like transcription factor.

### Ethephon Treatment Had No Effect on DAM Gene Expression

*DAM*s are considered to be the primary regulators of the dormancy cycle in peaches and other Prunus spp. (Castede et al. 2014; Zhu et al. 2020). Therefore, we examined the correlation between *DAM* expression and the regulation of PA/ACN biosynthesis in ET-treated and control trees. It was found that ET treatment did not have a significant effect on *DAM* expression (Fig. S12A), whereas *DAM2*, *DAM5*, and *DAM6* expression was significantly higher in the late blooming ‘KV021779’ cultivar than in the early-blooming ‘John Boy’ cultivar (Fig. S12B). The relative normalized expression of *DAM* genes showed higher induction at the beginning of endodormancy (200 and 400 CH), followed by a decrease to near the limit of detection at the end of dormancy in both treatments and genotypes. These results indicate that low temperatures in the fall lead to *DAM* upregulation to promote dormancy initiation, which may regulate ABA levels during dormancy-release periods to control PA/ACN biosynthesis.

## Discussion

This represented the first report of an association between two groups of marker metabolites (PA and ACN) and the release of endodormancy and ecodormancy, respectively, in peach floral buds. Analyses of metabolomics data, gene expression, PAs localization, and peach bloom time variations indicated that differential regulation of PAs/ACNs biosynthesis acted not only as a benchmark to define ET-mediated bloom delay but also a biomarker profile that distinguished early- and late-bloom cultivars of peach (Fig. 8). The findings showed that PA/ACN biosynthesis associated with bloom-related mechanisms occurred according to the activation of a gene module regulated by MYB transcription factors, ABA biosynthesis and ROS homeostasis. Identifying and characterizing these marker metabolites and their regulatory interactions could open new avenues for research in horticulture science related to dormancy regulation and should be expanded to other deciduous tree fruits.

**Figure 8.**
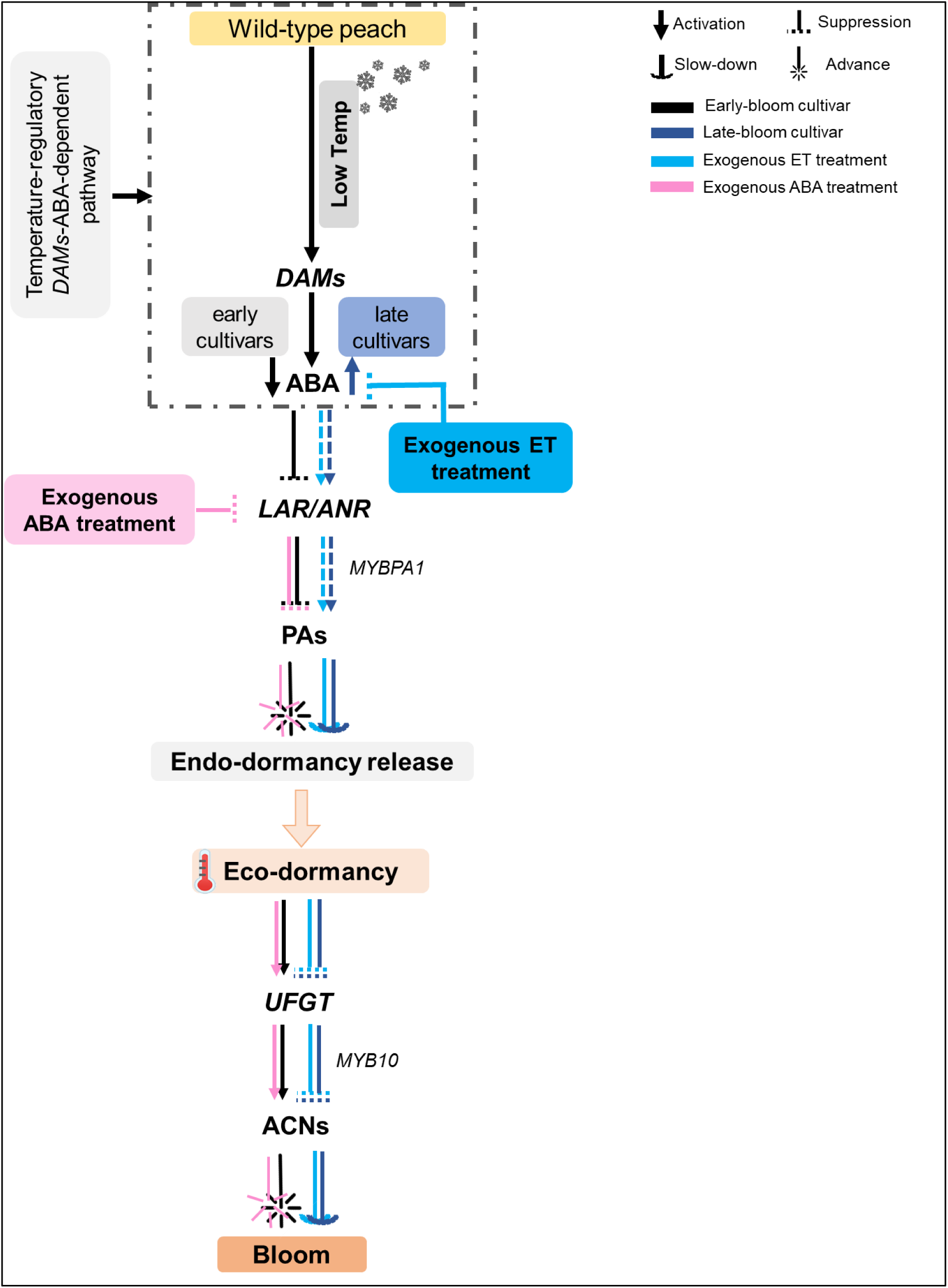
Model of the crucial role of PAs and ACNs in dormancy regulatory and non regulatory pathways for bloom time determination in peach. In dormancy regulatory peach genotypes, cold temperature positively upregulated *DAMs* leading to modulating ABA levels during endodormancy release. Higher ABA levels of early-bloom cultivar decreased PAs accumulation by suppressing *MYBPA1* transcriptional factor and *ANR/LAR* gene expression, leading to advanced endodormancy release. Progressive endodormancy release initiates ACNs biosynthesis by upregulating *MYB10* transcriptional factor and *UFGT* gene expression upon exposure to warm temperature during ecodormancy, consequently advancing bloom time. Exogenous ET treatment suppresses ABA levels during the endodormancy release, enhancing PAs accumulation to extend the dormancy release period, ultimately delaying bloom time. Exogenous ABA treatment downregulated the biosynthesis of PAs by suppressing *LAR*/*ANR* expression to advance bloom time. The black color indicates early-bloom cultivar, the deep blue indicates the late-bloom cultivar, the light blue indicates exogenous ET treatment, and the pink indicates exogenous ABA treatment.

Several lines of evidence presented suggest the significance of PAs accumulation during endodormancy and the association of ACNs accumulation at the end of ecodormancy with bloom time in peaches. Untargeted metabolomics data of peach floral buds revealed epicatechin as the only flavonoid strongly associated with endodormancy release among the 419 identified metabolites, with its level significantly higher following ET treatment compared to the untreated control (Fig. 2). During the endodormancy period, ACNs accumulation was almost undetectable, but it increased substantially along with ecodormancy progression, showing a 2-fold higher level in the untreated control compared to the ET-treated group (3000 GDH) (Fig. 4). Analysis of 12 peach genotypes with contrasting bloom dates showed significant variations in PA/ACN levels during the endodormancy and ecodormancy periods (Fig. 5). The PA/ACN ratio and DMAC staining of floral buds at 3000 GDH showed that late-bloom cultivars with high CRs contained significantly higher PA levels, which extended their endodormancy period, whereas floral buds of early-bloom cultivars had already reached their first-pink stage (Fig. 5, C and D). Previous studies on apricot (*P. armeniaca*) indicated that late-bloom apricot cultivars contain significantly higher flavonoids, including epicatechin and procyanidins, during the dormancy period (Conrad et al. 2019), supporting our findings. In contrast, in almond (*P. amygdalus*), the early-flowering phenotype was associated with a 3-fold increase in ACN content than late-flowering (Guillamon et al. 2020). The present study also showed a decline in PAs content in peach cultivars after endodormancy release that varied according to their respective CR (Table S1). Overall, these data support the notion that regulation of PA/ACN biosynthesis represents a signal for endodormancy release and bloom time in peaches.

ACNs and PAs are synthesized through a multi-step process from the common flavonoid biosynthetic pathway. The products of upstream biosynthetic genes such as PAL, CHS, CHI, and F3’H are involved in synthesizing common precursors (Shirley 2001). The cooperative action of late biosynthetic gene products catalyzes the downstream biosynthetic route to obtain ACN via the expression and translation of *DFR*, *ANS*, and *UFGT*. The translated products of *LAR* and *ANR* promote epicatechin synthesis, which leads to the production of polymeric PAs (Winkel 2001). MYB transcription factors such as PpMYB10 are activators of ACNs biosynthesis, whereas PpMYBPA1 is reportedly involved in regulating PAs biosynthesis in peach flowers (Zhou et al. 2016). In this study, *MYBPA1* and *ANR* expression was upregulated in the ET-treated group during endodormancy, whereas *MYB10* and *UFGT* expression was downregulated at the end of ecodormancy (3000 GDH). Furthermore, in the late-bloom cultivar (’KV021779’), higher PAs content along with upregulated *MYBPA1* and *ANR* expression during endodormancy were accompanied by lower transcript levels of *MYB10* and *UFGT* expression at 3000 GDH compared to the early-bloom cultivar (’John Boy’) (Fig. 5). These results provide additional evidence for a relationship between PAs/ACNs biosynthesis rate and bloom time regulation (Fig. 4).

In the present study, our findings indicate that the biosynthesis of PAs and ACNs during dormancy is likely regulated by ABA. First, untargeted metabolomics and target gene-expression analyses showed that ET treatment suppressed ABA levels during endodormancy (Fig. 6A). Second, the study of peach cultivars showed that ‘John Boy’, contained significantly higher ABA content throughout dormancy relative to that in the late-bloom cultivar, ‘KV021779’ (Fig. 6B). Third, a higher ABA level in the early-flowering cultivar was associated with suppressed PA biosynthesis, lower *ANR* expression, and advanced endodormancy release. In contrast, a lower ABA level in the late-flowering cultivar was combined with higher PA levels and delayed endodormancy release (Fig. 5). Fourth, the exogenous application of ABA during endodormancy (∼ 500 CH) advanced bloom time, reduced PA content and downregulated *LAR* expression in the floral buds of ‘Redhaven’ peach (Fig. 7). Together, our findings suggest that ABA-mediated suppression of PA biosynthesis during endodormancy might be necessary for the activation of ACN biosynthesis to initiate bud break and blooming. This inference is also in agreement with previous studies showing that ABA can modulate seed and bud dormancy by inhibiting *ANR*, *ANS*, and *DFR* expression (Guillamon et al. 2020; Zhao et al. 2019; Gao et al. 2018).

The decrease in ABA levels observed in peach trees during endodormancy following ET treatment can be attributed to the well-documented antagonistic relationship between ethylene and ABA (Arroyo et al. 2003; Lorenzo et al. 2003; Benschop et al. 2006; Ortega-Martinez et al. 2007; Dong et al. 2011). However, our observed interactions between ABA-PA and ethylene are particularly reminiscent of the ABA-ethylene relationship during stomatal opening and closure. It is well evident that ABA plays a critical role in regulating stomatal closure to mitigate the effects of abiotic stress, such as drought (Munemasa et al. 2015). However, exogenous application of ethylene inhibits ABA- and stress-induced stomatal closure in various plant species, including Arabidopsis, tomato, carnation, and sunflower (Madhavan et al. 1983; Tanaka et al. 2005). This inhibitory effect of ethylene on ABA-regulated stomatal activity results from the accumulation of flavonols, which is an EIN2-dependent process. EIN2 acts as a positive regulator of the pathway and is required for the downstream response to ethylene. Interestingly, flavonol accumulation represses ABA-induced ROS production and stomatal closure in Arabidopsis, particularly under stress conditions (Watkins et al. 2014).

Similarly, both control and ET-treated buds exhibited an increase in ABA levels (Fig. 6) following a significant decline in PAs levels observed at 800 CH during endodormancy (Fig. 5). The levels of H_2_O_2_ in the ET-treated buds also exhibited an earlier increase (at 600 CH) (Islam et al. 2021) that was inversely correlating with the decline of total PAs, epicatechin, and procyanidin B1 throughout the endodormancy period (Fig. 4). Notably, both PA and flavones originate from the same flavonoid pathway, with the former having an evident potent antioxidation capacity (Yu et al. 2020).

Taking all of this into consideration, it can be hypothesized that the shift in PA accumulation level in response to chilling accumulation during endodormancy is the driving force behind the increase in H_2_O_2_. This increase, in turn, induces ABA biosynthesis and subsequently ACNs during the ecodormancy period through bud break (Fig. 4–6). The synergistic interaction between ABA and ROS production has been confirmed in many examples, particularly during cold and other abiotic stresses. In some cases, ABA induces the production of ROS (Ibrahim and Jaafar 2011). However, a positive feedback mechanism has also been observed, in which the enhanced level of H_2_O_2_ caused by either drought treatment or exogenously supplied H_2_O_2_ activates the ABA biosynthesis gene *NCED3*, leading to ABA accumulation and promoting H_2_O_2_ generation (Park et al. 2021; Lee et al. 2022).

Dormancy-associated MADS-box (*DAM*) genes are directly influenced by chilling temperatures, and specific mutations in these genes have been associated with dormancy release or reduced chilling requirement, as reported by Yamane et al. (2011). During endo-dormancy, there is a self-enhancing regulatory loop between the ABA signal and *DAM*/SVP-like (SVL) proteins, whereby ABA promotes endo-dormancy and activates SVL expression. In turn, SVL activates *NCED3* expression, leading to increased ABA biosynthesis in dormant buds (Singh et al. 2018; Tylewicz et al. 2018). However, Tuan et al. (2017) have also reported a negative feedback regulation between ABA and *DAM*/SVP-like gene expression during bud dormancy.

The present study revealed that all six *DAM* genes exhibited peak expression levels at 200-400 CH, followed by a sharp decline, with no significant differences in gene expression between control and ET-treated trees. On the other hand, ABA levels increased steadily during endodormancy and peaked at the endodormancy release (1000 CH and 1000 GDH), with significant differences between control and ET-treated trees, indicating that these differences are not directly mediated by upstream *DAM* proteins. However, our study also observed differences in *DAM* expression between early- and late-blooming cultivars, with the latter showing higher transcript levels of *DAMs 2*, *5*, and *6* and lower ABA levels. Notably, in both model systems used in this study, ABA biosynthesis occurred after *DAM* genes showed a decline in their expression.

Based on our findings, and in conjunction with previous studies that have demonstrated higher levels of *DAM5* and *DAM6* expression in high-chill peach and apricot cultivars, as well as the observation that their expression levels reach a minimum at the same time as bud break competence acquisition in each cultivar (Jiménez et al. 2010), one could hypothesize that ABA accumulation begins earlier in low-chill/early-bloom cultivars. This hypothesis is based on the notion that the decline in *DAM* transcript levels initiates earlier in these cultivars. The early increase in ABA inhibits PA, enhances ACNs levels, and ultimately leads to early bloom. Although both *DAM* genes and total PA follow the same induction pattern during endodormancy (Fig. 4 and 5; Fig. S12), with peaks at 400 CH and declining thereafter, it is not yet clear whether this induction is mediated through ABA.

In summary, this study sheds light on the crucial roles of proanthocyanidins (PAs) and anthocyanins (ACNs) in regulating dormancy and bloom time in peaches. Specifically, epicatechin was identified as a key flavonoid associated with the release of endodormancy, and variations in PA/ACN ratios during dormancy were highly correlated with chill requirements and bloom time dates in 12 peach genotypes. Furthermore, the study reveals that ABA is a key regulator of the biosynthesis of PAs and ACNs during dormancy in peaches, as demonstrated by the suppression of ABA levels during endodormancy by ET treatment, and delayed endodormancy release in peach cultivars with higher ABA content throughout dormancy. Additionally, exogenous application of ABA during endodormancy was found to reduce PA content and advance bloom time. Moreover, differences in *DAM* expression were observed between early- and late-blooming cultivars, with the latter showing higher transcript levels of *DAM* genes and lower ABA levels. Finally, the study provides a possible model that could explain the link between *DAM* genes as the master genetic regulators of endodormancy and chill accumulation in stone fruits, and the downstream biochemical pathways that are mediated through the flavonoid pathway; particularly PAs and ACNs, ABA, and ROS.

Importantly, this study identifies PAs and ACNs as quantitative marker metabolites for endo- and ecodormancy, which would enable the development of more reliable chill requirement and heat requirement models. It is worth noting that the current models’ general invalidity and low transferability pose a challenge to accurately projecting the future consequences of global climate warming on the phenology of tree fruit species.

## Acknowledgment

We would like to thank Mr. Mark Demuth and Mr. Dennis Bennett from USDA-ARS, Appalachian Fruit Research Station, WV, USA, for assistance with peach bud sample collection.

## Author Contributions

**P.R.D.** contributed conceptualization, investigation, methodology, formal analysis, validation, writing–original draft preparation, visualization. **M.T.I. and J.L.** methodology, formal analysis, review and editing. **Z.L., and C.D.** methodology, review and editing. **S.M.S.** conceptualization, validation, resources, data curation, writing–original draft preparation; writing–review and editing, visualization, supervision, project administration, funding acquisition. All authors have read and agreed to the published version of the manuscript.

